# Thalamic volume and functional connectivity are associated with nicotine dependence severity and craving

**DOI:** 10.1101/2022.09.25.509385

**Authors:** Cindy Sumaly Lor, Amelie Haugg, Mengfan Zhang, Letitia M. Schneider, Marcus Herdener, Boris B. Quednow, Narly Golestani, Frank Scharnowski

## Abstract

Tobacco smoking is associated with deleterious health outcomes. Most smokers want to quit smoking, yet relapse rates are high. Understanding neural differences associated with tobacco use may help generate novel treatment options. Several animal studies have recently highlighted the central role of the thalamus in substance use disorders, but this research focus has been understudied in human smokers. Here, we investigated associations between structural and functional magnetic resonance imaging measures of the thalamus and its subnuclei to distinct smoking characteristics. We acquired anatomical scans of 32 smokers as well as functional resting-state scans before and after a cue-reactivity task. Thalamic functional connectivity was associated with craving and dependence severity, whereas the volume of the thalamus was associated with dependence severity only. Craving, which fluctuates rapidly, was best characterized by differences in brain function, whereas the rather persistent syndrome of dependence severity was associated with both brain structural differences and function. Our study supports the notion that functional versus structural measures tend to be associated with behavioral measures that evolve at faster versus slower temporal scales, respectively. It confirms the importance of the thalamus to understand mechanisms of addiction and highlights it as a potential target for brain-based interventions to support smoking cessation, such as brain stimulation and neurofeedback.

## Introduction

Nicotine use disorder (NUD) is associated with deleterious health outcomes and has been estimated to cause more than 6 million deaths per year worldwide (1). Most smokers want or attempt to quit but abstinence is notoriously difficult to maintain even when therapeutic strategies such as nicotine replacement therapy and nonnicotine medication were initially effective (2). Low cessation rates and high relapse rates emphasize the need to investigate factors responsible for the maintenance of NUD.

On a biological level, nicotine can fast cross the blood brain barrier, binds to nicotinic acetylcholine receptors (nAChR) and induces long-lasting cellular changes in dopaminergic pathways (3) that play a major role in reward processing and in the development and maintenance of addiction. The α4β2-nicotinic receptor, which has the highest affinity for nicotine, shows the highest density in the thalamus (4). Recent reviews have suggested that the thalamus acts as more than a relay station between the sensory system, the limbic system, and the cerebral cortex, and have highlighted the critical importance of the thalamus in substance use disorders (SUD) (5,6). Indeed, as a central part of cortico-striato-thalamo-cortical loops, the thalamus mediates reward processes (7), response inhibition (8) and salience attribution (9), which are central components in the development of SUD. In human neuroimaging studies, some cue-reactivity paradigms (10) revealed enhanced activation in the thalamus (as well as other brain regions) when smokers were exposed to smoking-related cues compared to neutral cues. Thalamic resting-state functional connectivity and volume have also been associated with smoking relapse (11).

On a psychological level, craving, or urge, is a core feature of NUD and a good predictor of cessation outcome (12). The urge to smoke can be provoked by external cues related to smoking such as watching someone smoke or smelling tobacco, but also by any other cue that the cigarette user has associated with smoking (e.g., drinking coffee, going to a bar, etc.). In a laboratory experiment, cue-induced cravings are often studied with cue-reactivity paradigms, which consist of measuring neural responses, bodily responses, or smoking urge levels when participants are exposed to various drug cues. However, removing external cues from a smoker does not mean that their urge to smoke disappears. Some internal “background” or baseline cravings persist after cue-exposure or spontaneously appear in the absence of any cue (13), as it can relate to states of stress and withdrawal. Although they are most likely not orthogonal dimensions, cue-reactivity measures do not necessarily correlate with baseline craving (13). A recent functional magnetic resonance imaging (MRI) study, which identified distinct insula-based neural substrates for smoking cue-reactivity measures and baseline craving (14), corroborates this observation.

While the NUD diagnosis is categorical, scales like the Fagerström Test for Nicotine Dependence (FTND) (15) add some dimensionality to the syndrome by characterizing its severity. Importantly, while NUD and its severity are rather stable constructs with time criteria explicitly defined by current diagnostic manuals (ICD/DSM), individual features defining NUD, such as craving, can, and do, fluctuate over time. Craving fluctuates within minutes or hours as a function of many factors that include nicotine blood level (16), exposure to smoking cues, and abstinence (17). On the other hand, the FTND assesses global attitude towards smoking and smoking habits, such as current daily cigarette consumption or propensity to smoke more in the morning compared to the evening, which is more stable than momentary craving measures. Test-retest reliability studies demonstrate high stability of FTND scores over weeks to months (18,19).

When looking for brain-behavior associations, it is important to consider the temporality of the chosen metric. Anatomical MRI scans allow estimating structural measures such as gyrification, white matter integrity, or subcortical volumes. Functional MRI (fMRI), in contrast, measures the blood-oxygen-level dependent (BOLD) signal, which is an indirect estimate of metabolic demands in localized brain regions. Although it seems possible to find some degree of interdependence between structural and functional measures (20), the temporal specificity of a brain measure could make it more suitable to capture facets of behavior that fluctuate at a similar temporal scale. Specifically, functional metrics, which tend to evolve faster than structural metrics, might be more suitable to capture state-like craving levels and structural metrics more suitable to capture trait-like dependence scores. Baur et al. (21), for instance, found that trait anxiety was more strongly linked to structural connectivity between the amygdala and the insula, while state anxiety was more linked to functional connectivity between the two regions. Similarly, Saviola et al. (22), found that trait anxiety mapped onto both structural and functional measures of the default mode network, while state anxiety mapped onto functional measures only. To our knowledge, no study so far has applied this state vs. trait distinction of tobacco smoking characteristics for brain-behavior associative studies.

To investigate the involvement of the thalamus in NUD, while keeping in mind that fast craving variations might be reflected in functional brain changes and more stable severity scores might be associated with more stable structural features, we analyzed resting-state data and anatomical data that were acquired together with our recent smoking cue-reactivity study (23). Since the thalamus is divided into subnuclei that carry different cognitive functions, we further fine-tuned our analysis by including thalamic subdivisions. While Haugg et al. (23) examined functional scans of the cue-reactivity task, here, we analyzed pre- and post-task resting-state scans, as well as high-resolution anatomical scans, in order to address the following hypotheses. First, we expect pre and post-exposure craving levels, as measured by the Brief Questionnaire of Smoking Urges (QSU) (24), to co-vary with respectively pre and post resting-state functional connectivity measures from thalamic seeds. We expect stronger brain-behavior associations during post-exposure than during pre-exposure since the cue-reactivity task increases craving levels and might harmonize mental states of the participants. Finally, we expect that NUD severity, as measured by the FTND, to correlate with thalamic volumes.

## Material and methods

### Participants

Participants were 32 individuals matching the DSM-5 criteria for NUD (age: 26.0±5.3; sex: 17F, 15M; daily cigarette consumption: 11.5±5.6, smoking history: 7.4±4.8 years of smoking, FTND score = 2.9±1.8). The FTND values of our sample ranged from 0 to 6 (mean = 2.6±1.8) on a scale of 0 (no dependence) to 10 (severe dependence). We chose not to exclude individuals with a null FTND score because correlational analyses benefit from a wider FTND distribution. Please note that it is possible to fulfill the DSM-5 criteria for NUD, while having no dependence as measured by the FTND. Although they show some overlap, the two questionnaires differ, with the FTND being generally more quantitative and more specific. Exclusion criteria were the use of non-cigarette tobacco substitutes such as nicotine patches, mental or neurological disorders and standard MRI-incompatibility (metal implants, pregnancy, etc.). We asked the participants not to smoke for at least one hour before the experiment.

The study was conducted in Zurich, Switzerland, in accordance with the Declaration of Helsinki, and was approved by the ethics committee of the University of Zurich. All participants gave their written informed consent after being informed orally and in writing of the eventual risks associated with MRI scans and the purpose of the study. They received a financial compensation (25CHF/h) for their participation.

### Experimental design

We exposed participants to a 20-minute-long cue-reactivity task, which consisted in passively watching 330 smoking-related or neutral pictures taken from image databases (e.g., the Smoking Cue Database (SmoCuDa) (25)). More details on the procedure and the stimuli can be found in (23). Immediately before and after viewing the smoking cues, we acquired resting-state scans of 7 minutes each (Figure 1). For the 7-min pre-task (pre-rest) and the po st-task (post-rest) resting-state runs, participants had to fixate a white fixation dot over a black background. A structural scan was acquired after the functional imaging acquisition. Before the scanning session, the participants filled out several behavioral questionnaires which included the Fagerström Test for Nicotine Dependence and the brief Questionnaire of Smoking Urges (QSU). The QSU was filled out a second time after the scanning session. Approximately 1 hour elapsed between the pre-QSU and the post-QSU. The data has been shared with the ENIGMA Consortium (https://enigma.ini.usc.edu; https://www.enigmaaddictionconsortium.com/). The data will also be made available by the authors upon reasonable request.

**FIGURE 1.**
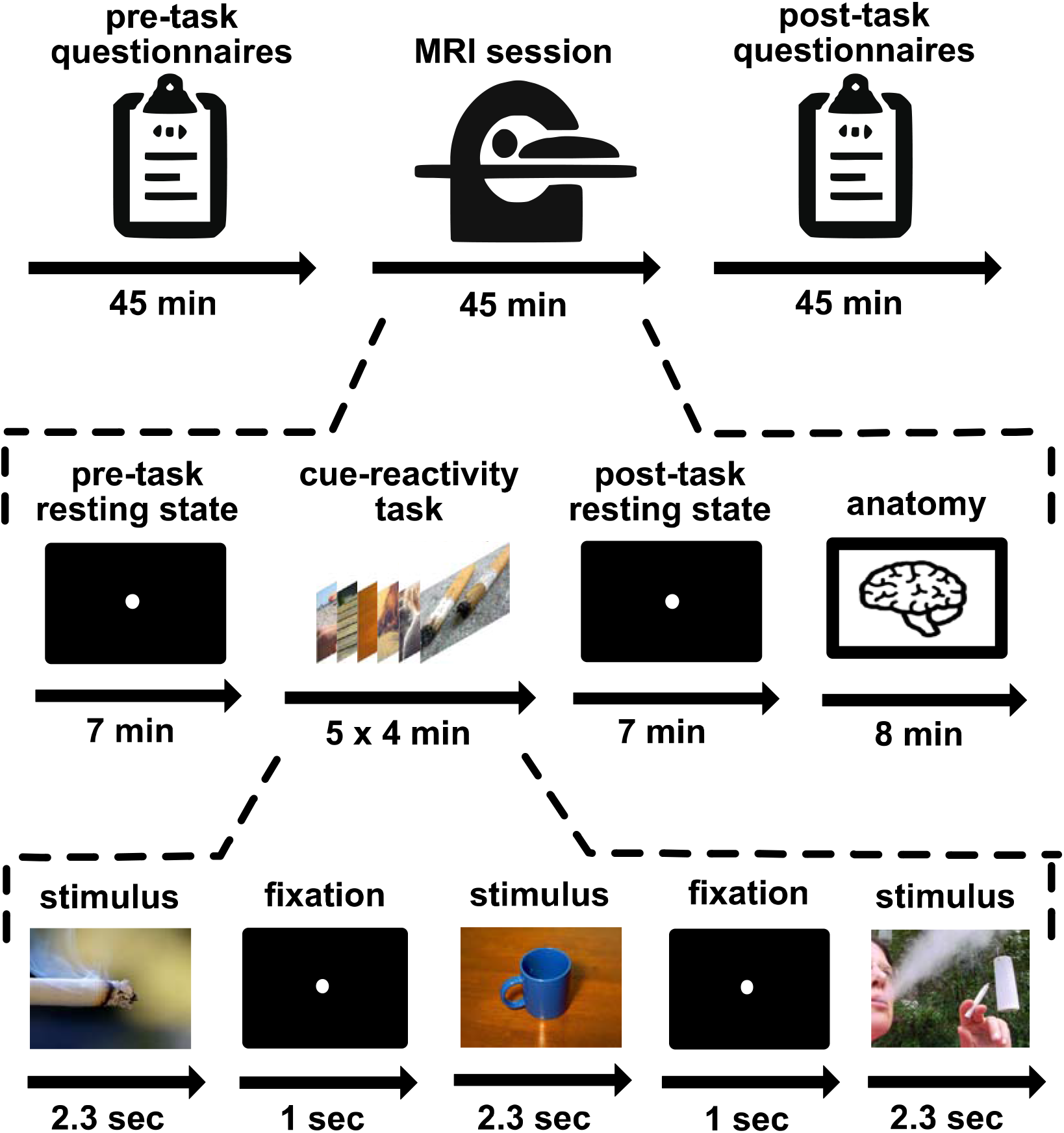
Experimental design. The study was divided into three parts. In the first part, subjects filled out several questionnaires on their smoking routines. Then, subjects underwent 50 min of scanning, including resting state scans, five runs of passive viewing and an anatomical scan. During each passive viewing run, 68 out of a total of 340 neutral or nicotine-related images (including 10 additional catch trial images) were presented for 2.3 s in random order, followed by a 1-s fixation dot baseline. Finally, subjects rated all 330 presented images with respect to craving and valence. Reprinted with permission from Wiley & Sons Ltd (Haugg et al., 2021).

### MRI acquisition

MR data was collected with a 3.0 Tesla Philips Achieva system (Philips Healthcare, Best, The Netherlands) using a 32 phased-array head coil at the Psychiatric University Hospital, Zurich. The two resting-state functional scans were acquired with a T2*-weighted gradient-echo planar imaging (EPI) sequence (repetition time (TR) = 2000ms, echo time (TE) = 35ms, flip angle (FA) = 82°, 33 slices in transverse orientation, no slice gap, voxel size = 3 x 3 x 3 mm^3^, field of view (FoV) = 240 x 240 x 99 mm^3^, total scan duration = 7:12 min) and. At the end of the session, we additionally collected a high-resolution anatomical T1-weighted scan (FA = 8°, 237 slices in sagittal orientation and ascending order, no slice gap, voxel size = 0.76 x 0.76 x 0.76 mm^3^, FoV = 255 x 255 x 180 mm^3^, total scan duration = 08:26 min).

### Thalamic segmentation

To estimate the volume of the thalamus from anatomical data, we first visually inspected the raw anatomical scans to ensure data quality. In two participants, we identified wave-like artifacts, which we corrected with the spatial-adaptive Non-Local Means (SANLM) denoising filter of the CAT12 toolbox (http://www.neuro.unijena.de/cat/). Next, we performed subcortical segmentation using FreeSurfer software V7.1.1 (https://surfer.nmr.mgh.harvard.edu/). The thalamic volume and the cortical volume of left and right hemispheres were calculated following the standard segmentation procedure in Freesurfer (http://surfer.nmr.mgh.harvard.edu/fswiki/recon-all) which performs cortical reconstruction and volumetric segmentations. This fully automated segmentation and labeling approach consists of five stages (1. affine registration to MNI305 space; 2. initial volumetric labeling; 3. correction of intensity variation due to B1 bias field; 4. alignment to MNI305 atlas; 5. volume labeling; https://surfer.nmr.mgh.harvard.edu/fswiki/FreeSurferAnalysisPipelineOverview). Each participant’s image was later visually examined for misalignment or distortions, and no further manual corrections were added.

As a second step, we applied Freesurfer’s procedure for thalamic segmentation into different nuclei (26) (https://freesurfer.net/fswiki/ThalamicNuclei) to obtain the volume of the nuclei for each participant. For the sake of consistency, we included the same nuclei as in the functional connectivity analysis (described below), i.e., the anteroventral nucleus (AV), Lateral Posterior (LP), Ventral Anterior (VA), Ventral Lateral (VL), Ventral posterolateral (VPL), Intralaminar (IL), MedioDorsal (MD), Lateral geniculate (LGN), Medial geniculate (MGN), Pulvinar (Pu). Since the procedure parcellates the MD, the Pu, the VL and IL into substructures by default, we later combined them into one region each. We did not include the Nucleus Reuniens due to its small size (only 2 voxels per hemisphere for functional data).

### Seed-based functional connectivity analysis

We used MATLAB2018a and the CONN20b toolbox (www.nitrc.org/projects/conn, RRID:SCR_009550) to analyze functional scans. We preprocessed neuroimaging data using CONN’s default preprocessing pipeline (which included functional realignment and unwarp, slice-timing correction, ART-outlier identification, segmentation and normalization, and functional smoothing with an isometric 6 mm Gaussian kernel) and a standard denoising pipeline (band-pass filtering (0.008-0.09 Hz), linear de-trending, aCompCor to remove confounds by specifying realignment parameters (27), ART-derived outliers and the first five principal components of white matter masks and cerebrospinal fluid masks as nuisance covariates). Three participants were excluded from functional connectivity analyses because of high motion, which we defined as mean framewise displacement (28) above 0.3 mm or less than 5 min of valid scans remaining after ART-outlier scan censoring in one or more resting-state runs.

We used a seed-based analysis with the left thalamus and the right thalamus from the Harvard-Oxford atlas (https://fsl.fmrib.ox.ac.uk/fsl/fslwiki/Atlases) as seed regions. For each seed, each run, and each participant, Fisher Z-transformed brain maps were created by calculating the correlation coefficient between the timeseries of the seed region (timeseries averaged across all the voxels in the seed) and the timeseries of each other voxel. The same analysis was then repeated using each subnuclear division of the thalamus as the seed. The masks (AV, LP, VA, VL, VPL, IL, MD, LGN, MGN, and Pu, left and right) were taken from the AAL3 atlas, which is based on Freesurfer’s thalamus segmentation method (29).

### Test for brain-behavior associations

To test for statistical associations between FTND scores and thalamic volumes (left and right), we used Spearman partial correlations (MATLAB2018a) for which we specified age, sex, as well as the cortical volume of the corresponding hemisphere as co-variates of no interest. Since we tested 11 regions for each hemisphere, we applied the Bonferroni correction to control for multiple testing (22 tests in total). We additionally tested whether thalamic volumes were associated with smoking urge scores (i.e., scores on the QSU) using the same partial correlations between urge to smoke scores (pre and post) and 22 thalamic volumes (left and right) and applied Bonferroni correction.

To test for statistical associations between smoking urge scores and resting-state functional connectivity, we performed a regression analysis on functional connectivity maps with smoking urge scores as a regressor of interest for each run (pre and post) and each thalamus seed (right and left). Specifically, we used pre-task urge scores as a regressor on pre-task brain maps and post-task urge scores as a regressor on post-task brain maps. One participant did not complete the QSU and was excluded from the functional connectivity analyses. Standard thresholding criterion based on Random Field Theory (30) was applied to select for significant clusters while correcting for family wise errors (voxel-level p_-uncorrected_ < 0.001 and cluster-level p_-FWE_ < 0.05). Since we performed 22 tests, we applied an additional Bonferroni correction by lowering the significance threshold for clusters to p_-FWE_ = 0.05/22 = 0.00113. Significant clusters were labelled with Automated Anatomical Labeling (AAL) atlases (31) as implemented in CONN20b.

An additional analysis was then performed using FTND scores instead of urge scores as regressor of interest, for pre- and for post-exposure task resting state runs, and each seed, and with Bonferroni correction.

## Results

### Behavioral data description

A paired t-test showed that smoking urge scores, as measured with the QSU (pre-urge = 10.0±30.7; post-urge = 46.6±30.0) significantly increased after the passive viewing task (p < 0.0001), the value being multiplied by more than a factor of 4 on average. FTND scores were positively correlated with pre-urge and post-urge scores, albeit not significantly (pre-urge: r = 0.32, p = 0.072; post-urge: r = 0.342, p = 0.060).

### Positive correlation between thalamic volumes and nicotine dependence

FTND values were significantly positively correlated with the volume of the whole right thalamus: (□ =0.056, p-corrected = 0.034, p_-uncorrected_ = 0.0016) (Figure 2). Of note, the left thalamic volume (□ = 0.49, p_-corrected_ = 0.16, p_-uncorrected_ = 0.0074), as well as the left IL nucleus (□ = 0.45, p_-corrected_ = 0.32, p_-uncorrected_ = 0.014) and the left MD nucleus (□ = 0.40, p_-corrected_ = 0.67, p_-uncorrected_ = 0.030) were also positively correlated with FTDN values (p_-uncorrected_ < 0.05), but the results did not survive Bonferroni correction.

**FIGURE 2.**
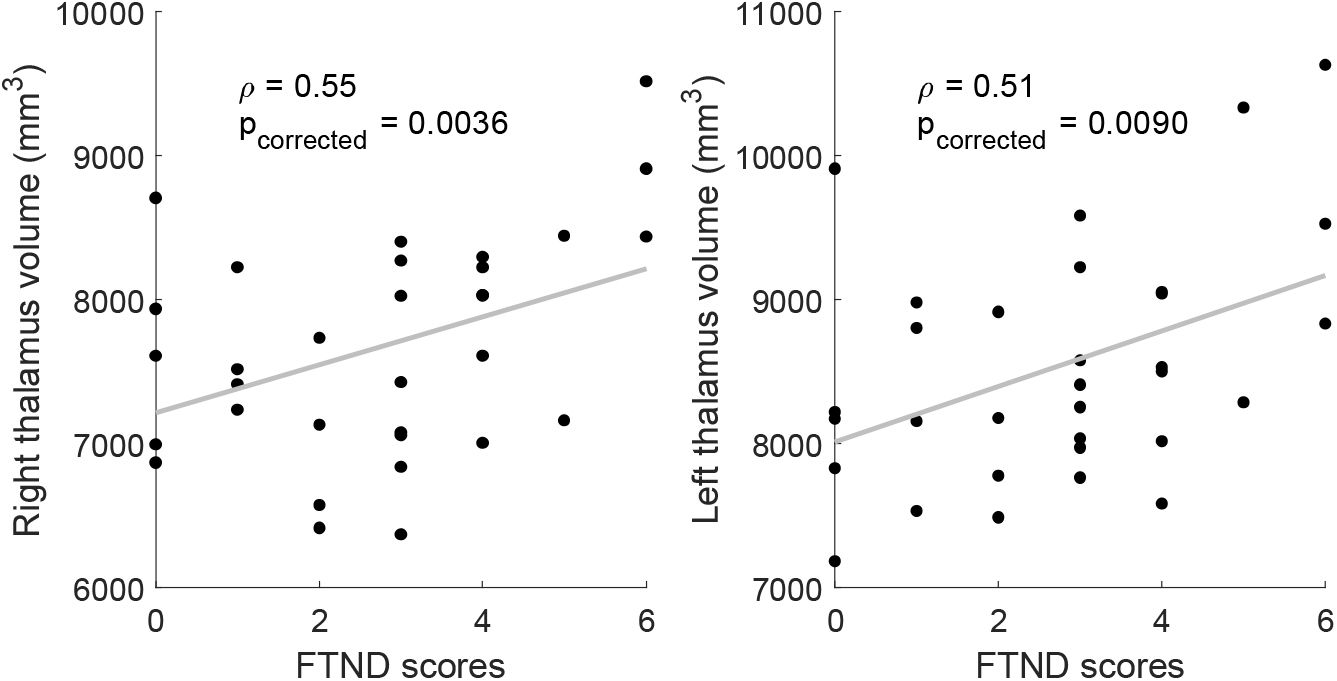
Thalamic volumes were positively correlated with Fagerström nicotine dependence scores. Thalamic volume and total intracranial brain volume (eTIV) were estimated using Freesurfer automatic segmentation of subcortical areas. Spearman partial correlations between nicotine dependence and thalamic volume (left and right) were corrected for age, sex and brain volume, and we applied the Holm-Bonferroni to correct for multiple testing. N = 32.

We found no significant association between the volume of the thalamus or its subnuclei (left or right) and urge scores (pre or post).

Please note that the thalamus subdivision using 3T scans have very recently been shown to not always be robust (32). Thus, these results should be interpreted with caution.

### Associations between thalamic functional connectivity and smoking urge

Regression analyses between urge scores and whole thalamus-based connectivity maps revealed that the functional connectivity between the whole left thalamus and a cluster in the dorsal part of the anterior cingulate cortex (ACC) (MNI peak coordinates (x, y, z) = (+6, +10, +32); size = 150 voxels; cluster p_-FWE_ = 0.000477) negatively co-varied with urge scores after task exposure (Figure 3) but not before.

**FIGURE 3.**
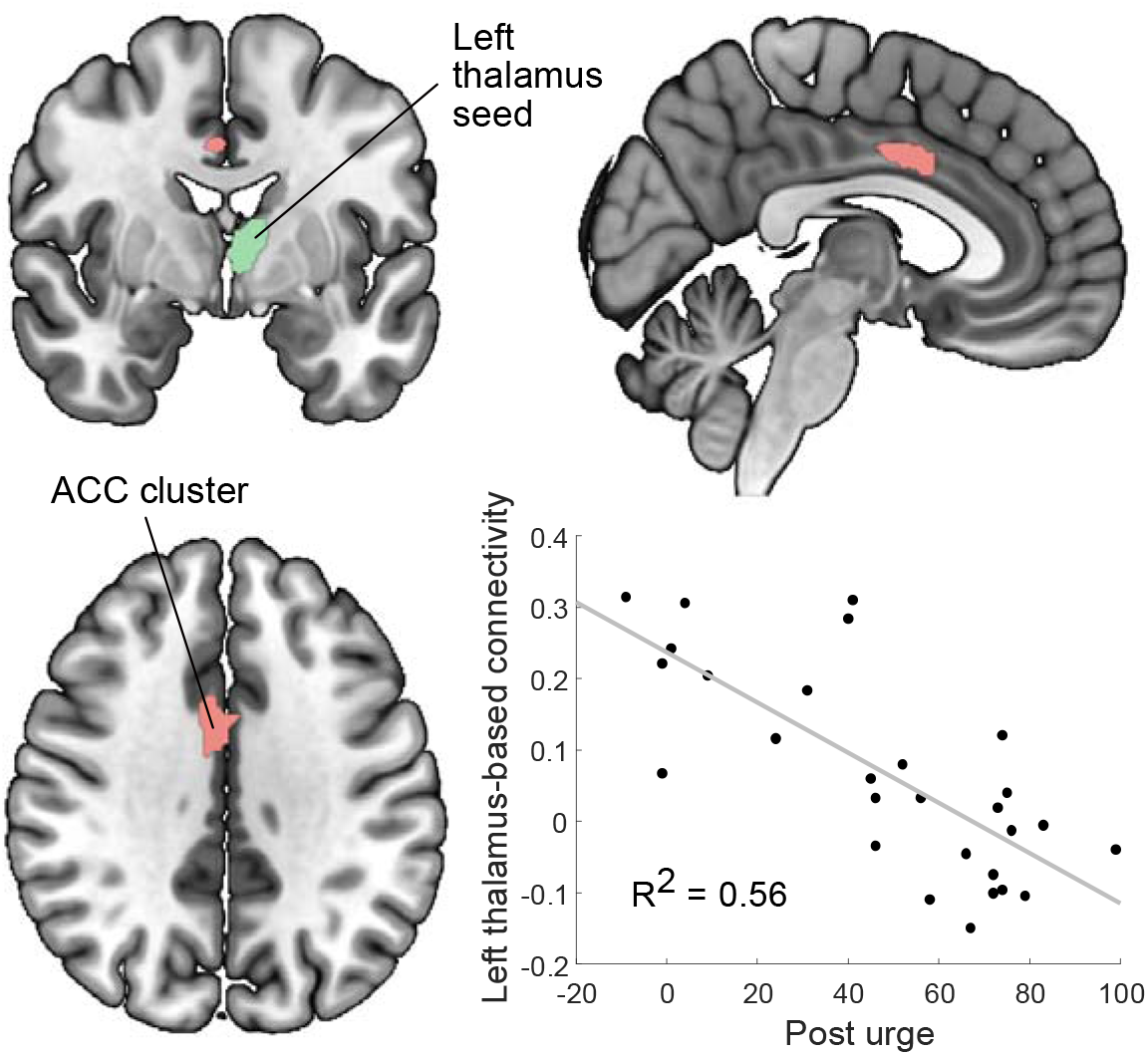
Regression analyses applied on thalamus-based correlation maps, with urge to smoke scores as the regressor of interest. Left thalamus-based functional connectivity to a cluster in the anterior cingulate cortex (ACC) co-varied with urge scores in the post-task run (MNI peak coordinates = +6, +10, +32; size = 150 voxels; p-FWE = 0.000477 < 0.05/4). Functional connectivity values between the left thalamus and the ACC cluster were plotted against urge scores (R^2^ = 0.56198) for illustrative purposes. N = 28.

Further, by dividing the thalamus into subnuclei as seed masks, a regression analysis with post-urge scores revealed a negative association with left VL-bilateral insula rsFC as well as with right Pu-ACC rsFC, and a positive association with right Pu-dorsolateral prefrontal cortex rsFC. The pre-urge-based regression analysis revealed a positive association with left LP-supramarginal gyrus rsFC.

Regression analyses with urge scores and other subnuclei as seeds did not reveal any other significant cluster. However, the same analyses using the pre-rest functional connectivity maps and thalamic seeds but with the FTND as regressor of interest identified a Right IL-Lingual gyrus/Precuneus and a Right MNG-Middle temporal gyrus association. Detailed results are reported in table 1.

**Table 1.** Seed-based functional connectivity results. Clusters were labelled using the Automated Anatomical Labeling (AAL) Atlas by the CONN toolbox. The table reports all clusters with p-uncorrected < 0.001 at the voxel level and p-FWE < 0.05/22 at the cluster level. Abbreviations: FC: functional connectivity; AV: anteroventral, LP: lateral posterior, VA: ventral anterior, VL: ventral lateral, VLP: ventral posterolateral, IL: intralaminar, MD: mediodorsal, LGN: lateral geniculate, MGN: medial geniculate, Pu: pulvinar nucleus.

| **Regression variables** | **Seed** | **Peak MNI coordinate** | **Size** | **Cluster p-FWE** | **Corresponding region** | **+/- association** |
| --- | --- | --- | --- | --- | --- | --- |
| Pre-rest FC<br>Fagerstroem | Right IL | -16 -58 +00 | 174 | 0.000044 | Lingual gyrus left<br>Precuneus | - |
|  | Right MNG | -48 -22 -12 | 210 | 0.000004 | Middle temporal gyrus | - |
| Pre-rest FC<br>Pre-Urge | Left LP | +44 -40 +44 | 164 | 0.000066 | Supramarginal gyrus | - |
| Post-rest FC<br>Post-Urge | Left VL | -30 +12 +08 | 259 | 0.000001 | Insula Left | - |
|  |  | +42 +14 +14 | 207 | 0.000015 | Insula right | - |
|  | Right Pu | -42 +50 +10 | 180 | 0.000087 | Dorsolateral prefrontal<br>cortex | + |
|  |  | +10 +06 +42 | 157 | 0.000276 | ACC | - |
|  | Left Whole<br>thalamus | +6 +10 +32 | 150 | 0.000477 | ACC (dorsal) | - |

## Discussion

The thalamus is a central node of the cortico-striato-thalamo-cortical reward circuit but has been understudied in drug addiction research compared to striatal regions (5). Here, we report associations between smoking urge and brain connectivity as well as thalamic brain volume and NUD severity, respectively. First, in structural data, we found that the volume of the thalamus positively correlated with NUD severity. Second, in resting-state data, using seed-based functional connectivity with the thalamus as a seed, we found a associations between smoking post-task urge scores and the connectivity between the whole left thalamus and a cluster in the dorsal ACC, between the right Pu and clusters in the dlPFC and the ACC, and between the left VL and bilateral insula. Our data seems to support the idea that state-like smoking urge was associated with functional measures, whereas trait-like NUD severity was not only associated with anatomical measures but also with functional measures.

Some molecular effects of nicotine could explain why individuals with severe NUD might have acquired a larger thalamus compared to individuals with low severity. Smoking is known to promote the upregulation of nAChR, which are found with largest densities in the thalamus (33). Also, higher levels of nAChR before smoking cessation treatment increases the likelihood of treatment success (34). Interestingly, many genetic polymorphisms of nAChR have been directly linked to NUD severity (35). Early attentional processes such as sensorimotor gating, which is involved in craving (36) and linked to the nicotine effects on the thalamus (37), are also connected to polymorphisms of the nAChR α3-subunit (38) known to be involved in the genetic risk for smoking (39). To the best of our knowledge, the effects of these polymorphisms on the function of the receptors, specifically whether these polymorphisms favor more upregulation, have not been formally characterized yet. Nevertheless, current evidence is compatible with the following speculations: a) individuals with higher NUD severity carry genetic predispositions for higher nAChR upregulation, and b) nAChR upregulation promotes thalamic volume increase. To support this last hypothesis, Positron Emission Tomography (PET) studies in rodents and humans (40,41) linked exposure to nicotine to greater nAChR densities in many brain regions with the exception of the thalamus. One possible explanation is that the high nAChR density in the thalamus left no molecular ‘space’ for additional receptors, which could imply that the thalamus itself would have to expand to accommodate the new receptors. Such structural expansion would not occur in other brain regions, where density had not yet reached a point of saturation. Moreover, refining our results with the analysis of thalamic subnuclei allowed to further support our suppositions. A recent study (42) found that the MD and the centromedian (CM) nucleus show the highest nAChR density, suggesting that these regions would mostly drive the volumetric increase. This is in line with our results, as here, the MD and the IL nuclei (which includes the CM) were the nuclei that correlate the most with NUD severity. To our knowledge, however, a causal relationship between nAChR upregulation and thalamic volume increase is still hypothetical and needs further investigation.

Strikingly, the direction of correlation between thalamus volume and smoking behavior is inconsistent in the literature. Some studies did not find significant volume differences between smokers and non-smokers (43). In more recent studies that also used Freesurfer for segmentation, operationalizations of worse smoking status (e.g. smoker > nonsmoker, high FTND > low FTND, relapser > non-relapser) were sometimes associated with larger volumes, but sometimes smaller volumes. Moreover, apparent contradictions can be found within the same study. For instance, Wang et al. (11) found larger thalamus volumes when contrasting smokers to age-matched non-smokers and relapsers to non-relapsers, but a negative correlation between the left thalamic volume and FTND scores. Yu et al. (44) also found a negative correlation between volume and FTND, and smaller volume when contrasting smokers and nonsmokers. In our sample, the correlation between volume and FTND was positive for both the left and right volumes. Apart from minor methodological differences (we corrected the left and right thalamic volume with cortical volumes of the corresponding hemisphere whereas they corrected with total intracranial volumes), inconsistencies could be explained by large differences in demographics. For example, Yu et al. (42) only included very young participants (16 to 24 y.o.) who mostly started smoking when they were teenagers (15 y.o.). By contrast, Wang et al. (11) and our study included older participants who started smoking when they were young adults. This is of critical importance since smoking is thought to interfere with the structural development of the brain (45), which could explain our opposite results. According to Wang et al. (11), it could also be that young individuals with smaller thalamus would be more likely to start smoking, but exposure to nicotine would increase thalamic volume in such a way that they have larger thalamus compared to age-matched non-smokers once they are older. To investigate potential effects of smoking history (years of smoking), we ran a post-hoc FTND/volumes correlation analysis that also includes the number of years smoking as a control variable (in addition to sex, GM volume and age). Further, we correlated thalamic volumes with years smoking. Interestingly, both analyses found minimal to no influence of smoking years on the FTND/right volume association, and no other volume correlated with years (detailed results in supplementary material). This further supports that smoking-related brain adaptations manifest in a multifactorial way that might include a genetic component. Divergences in the literature can also be explained by the fact that the three studies recruited samples with different ranges of FTND scores. The FTND scores in our sample were lower (2.9±1.8) than those of the two previous studies (5.0±1.9; 5.2 ± 2.2), and unlike them, we included individuals who, despite qualifying as smokers, were not dependent. They found a moderate negative FTND – thalamus volume correlation whereas we found a positive one. Also, Wang et al. (11), despite finding a negative correlation, found lower volumes for non-smokers, which, at first glance, seems more consistent with our results than theirs. However, this apparent contradiction can be explained by the fact that Wang’s negative correlation seems mostly driven by high FTND individuals. Combined, our studies seem to be compatible with FTND showing a general non-monotonic association with the volume of the thalamus (inverted U-shape with a peak occurring around FTND = 6). This highlights the complexity of brain-behavior associations and the need to refine our understanding of mechanisms of brain adaptation to smoking. To understand trajectories of smoking-induced brain changes, more diverse and larger samples are needed, as well as more within-subject longitudinal data. Indeed, prospective studies that track participants’ structural and smoking features before and after the onset of NUD would be necessary to understand whether smoking causes thalamus enlargement and whether a larger volume is a predisposition to higher severity, these not being mutually exclusive.

Additional findings were associations between smoking urge and thalamic functional connectivity during posttask rest, as well as between NUD severity and pre-task thalamic rsFC. In line with this, Saviola et al. (22), who mapped trait vs. state anxiety to brain measures, also found that trait mapped to both functional and structural data, while state only mapped to functional data. Here, we notably found that post-urge co-varied with the rsFC between (a) the whole left thalamus and the dorsal ACC, (b) the left VL and bilateral anterior insula, and (c) the right Pu and clusters in the ACC and the dlPFC. The ACC, the dlPFC and the insula have repeatedly been shown to be associated with craving in smokers. Their involvement in SUD-related processes has motivated neurofeedback researchers to train smokers to voluntarily control the activity levels of these regions as a mean to support smoking cessation (46–49). Interestingly, one of these neurofeedback intervention studies trained these exact three regions (49).

Craving induced by abstinence from smoking has previously been linked to connectivity between the left thalamus and a cluster in the right ACC (50), and might suggest similar neural substrates for both cue-induced craving and abstinence-induced craving. Besides, the VL nucleus is an integrative center for motor control (51) thus its connectivity with bilateral insula might be linked to transfer of interoceptive awareness to motor regions. The connection between the Pu and the ACC and dlPFC might be related to selective visual attention to smoking cues (52). In all, our functional connectivity results tend to confirm that the thalamus serves as a functional hub that mediates several cognitive components of SUD such as craving, motor planning, interoceptive awareness and visual attention.

### Limitations

This study carries several limitations. First, we did not formally assess whether the participants complied to the instruction of not smoking for one hour prior to the experiment. However, all participants confirmed having adhered to these instructions and we do not have reasons to doubt their compliance. Even if a few participants smoked just before MR-scanning, it is unlikely that this confound can explain the systematic and robust effects that we found. Second, we cannot definitely determine whether a larger thalamus precedes smoking, whether smoking causes the enlargement of the thalamus, and whether enlargement is modulated by genetic dispositions linked to NUD severity. To answer these questions, high sample size longitudinal studies which track the participants before and after the onset of smoking and their precise smoking patterns will be required. The lack of, for example, Positron Emission Tomography scans of nAchR, genetic data (e.g., on nAchR polymorphisms), as well as unclear molecular mechanisms (e.g., effect of NUD severity – related nAchR polymorphisms on kinetics of upregulation, and effect of upregulation on volumetric enlargement) prevent a detailed molecular-level interpretation of our findings. Another limitation along the same lines is the lack of a non-smokers control group for both the functional and the structural data, as well as a non-smoking-related task as a control condition for the functional study. Therefore, we do not know whether our connectivity and structural values fall under normal range or not.

### Conclusion

To summarize, this study distinguished between the mapping of state-like urge to smoke scores and thalamic-ACC resting-state connectivity, versus between thalamic volumes and trait-like dependence scores. Our findings suggest that the thalamus is a strong candidate for further multimodal and longitudinal investigations on the neural basis of different characteristics of NUD. Eventually, a better understanding of the neural underpinnings of NUD and craving could facilitate the development of novel interventions to support smoking cessation, such as neurofeedback, which requires determining precise targets and choosing temporally suitable neuroimaging and behavioral outcome measures.

