## Supplementary material for "Thalamic volume and functional connectivity are associated with nicotine dependence severity and craving"

We ran supplementary analyses to rule out potential effects of smoking history (number of years smoking) on our thalamic volumetric analyses.

We first correlated years of smoking with pre-urge scores (r = -0.246, p = 0.183), post-urge scores (r = 0.020, p = 0.914), and FTND scores (0.134, p = 0.464) and found no association.

Next, we verified that years of smoking did not influence the association between volumes and FTND by adding years of smoking as another control variable (in addiction to age, sex and hemispheric grey matter volumes) of our partial analyses.

We found that this control variable marginally impacted the results. FTND values were still significantly positively correlated with the volume of the whole right thalamus: (ϱ =0.055, p_-corrected_ = 0.050, p_-uncorrected_ = 0.0023). Likewise, the left thalamic volume (ϱ = 0.48, p_-corrected_ = 0.20, p_-uncorrected_ = 0.0090), as well as the left IL nucleus (ϱ = 0.46, p_-corrected_ = 0.32, p_-uncorrected_ = 0.015) and the left MD nucleus (ϱ = 0.39, p_-corrected_ = 0.91, p_-uncorrected_ = 0.041) were also positively correlated with FTDN values (p_-uncorrected_ < 0.05), and the results did not survive Bonferroni correction.

Similarly, we found no significant association between thalamic the volume of the thalamus and its subnuclei (left or right) and urge scores (pre or post).

Finally, we applied the partial correlation analysis between years of smoking and all thalamic volumes, controlling for age, sex and hemispheric grey matter volume, but found no significant associations, with or without Bonferroni correction.
